# Ancient DNAs and the Neolithic Chinese super-grandfather Y haplotypes

**DOI:** 10.1101/487918

**Authors:** Ye Zhang, Xiaoyun Lei, Hongyao Chen, Hui Zhou, Shi Huang

## Abstract

Previous studies identified 3 Neolithic Han Chinese super-grandfather Y haplotypes, O2a2b1a1a-F5, O2a2b1a2a1-F46, and O2a1b1a1a1a-F11, but their relationships with the archaeological and written records remain unexplored. We here report genome wide DNA data for 12 ancient samples (0.02x-1.28x) from China ranging from 6500 to 2500 years before present (YBP). They belonged to 4 different genetic groups, designated as Dashanqian (DSQ) of Xiajiadian Culture in the Northeast, Banpo (BP) of middle Yangshao Culture in the Central West, Zhengzhou Xishan (ZX) of Miaodigou Culture in the Central Plains, and Others. Present day F5 samples were closer in autosomal distances to the ZX and DSQ groups while F11, C, O1, and O2 samples were closer to the BP group. We also sequenced the Y chromosome of one of these ancient samples K12 from DSQ and found both K12 and a previously reported ~4000 year old sample MG48 from Northwest China to have the O2a2b1a1a1a2a-F2137 haplotype, belonging to the most prolific branch O2a2b1a1a1-F438 immediately under F5. We further found close relationships between ZX and DSQ and between ZX and ancient M117 Tibetans or present day Southwest Dai Chinese carrying the F5 subtype O2a2b1a1a6, implicating radiations of F5 subtypes from the putative place of F5 origin in ZX. These results are remarkably consistent with archaeological and written records.

## Introduction

There are numerous data for human activity in China from the time of the Neolithic period to the beginning of written records. There were the Gaomiao and Pengtoushan Culture of ~5800 BC in the South (Hunan), the Jiahu Culture and Peiligang Culture of 7000-5000 BC in the Central Plains (Henan), the Xinglongwa Culture of 6200-5400 BC and later the Hongshan and Xiajiadian Cultures in the Northeast (Inner Mongolia-Liaoning border), the Dadiwan Culture of 5800-5400 BC in Gansu and Western Shaanxi ^1^. At 5000 BC to 3000 BC, the Yangshao Culture was the most popular and existed extensively along the Yellow River in China and flourished mainly in the provinces of Henan, Shaanxi and Shanxi with the early period of the Culture mostly found in Shaanxi and the late period in Henan ^1^. Elements of the middle to late Yangshao Culture, the Miaodigou Culture, have been found widely in China, including the Hongshan Culture in the Northeast (3500 BC), indicating the broad cultural migration and influence of this Culture ^2^.

By analyzing the Y chromosome haplotype patterns, three Neolithic super-grandfather haplotypes have been discovered that together account for ~40% of present day Han Chinese males ^3–5^. The expansion dates are estimated 5400 YBP (years before present) for O3a2c1a-M117-F5 (O2a2b1a1a-F5 or F8, ISOGG 2017), 6500 YBP for O3a2c1-F46 (O2a2b1a2a1-F46), or 6800 YBP for O3a1c-002611-F11 (O2a1b1a1a1a-F11), and these three haplotypes represent 16%, 11%, and 14% of present day Han Chinese males, respectively. Several historical writings on ancient Chinese mention the great leaders/ancestors around the time of 5000 years ago or earlier, including *Fu Xi*, *Yan* Emperor (Yandi), *Huang* Emperor (Huangdi), and *Chi You*.

It remains unclear how the Neolithic Cultures were related to the super-grandfather haplotypes and how the different Neolithic Cultures were interconnected. We here addressed these questions by analyzing ancient DNA samples from 10 different sites in Central and Northern China. We found evidence of F5 associated autosomes in the Miaodigou Culture in Henan, F11 associated autosomes in the early Yangshao Culture in Banpo, and both F5 haplotype and F5 associated autosomes in the Xiajiadian Culture in Inner Mongolia. The results provide a coherent account of information from the relevant fields.

## Results

### Relationships among aDNAs in Central and Northern China

We collected 12 skeletal remains from 10 sites in Central and Northern China that were 2500-6500 years old (Table 1). DNAs were extracted and sequenced to different degrees of coverage (0.018x-1.27x). SNPs were called using the Hg19 reference genome. We obtained two sets of SNPs, one slow and the other fast, with the slow set informative for phylogenetics and the fast set representing natural selection and maximum saturation as detailed before ^6–12^. We then merged the SNPs of each of the aDNA samples with those of the 1000 genomes project (1kGP) ^13^, and obtained pairwise genetic distances, i.e., mismatch in homozygous (hom) sites for slow SNPs or mismatch in all sites for fast SNPs, between each aDNA sample and all individuals in the 1kGP.

**Table 1.**
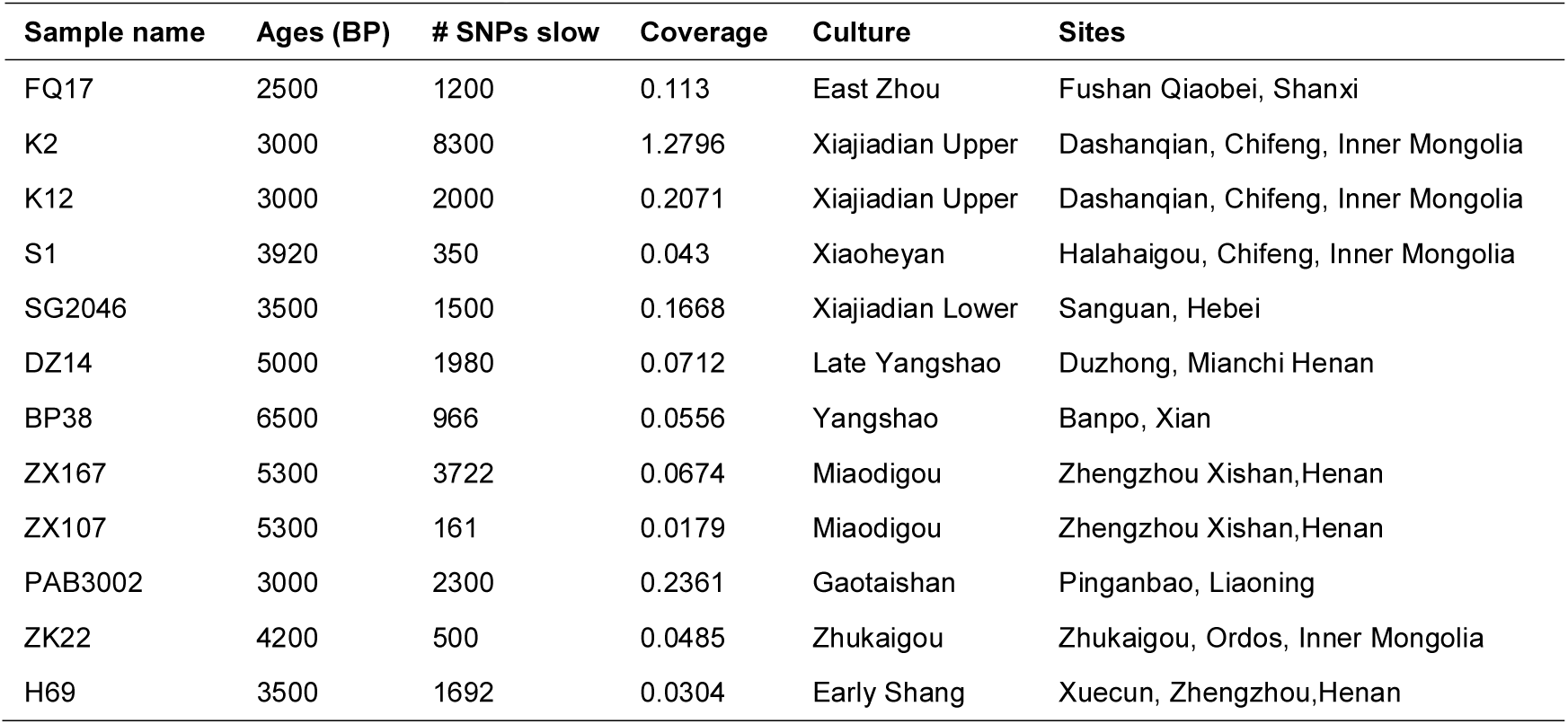
Information on ancient samples for which we report the nuclear sequence data in this study.

Because different ancient samples were sequenced at different coverages, it is unrealistic to use SNP mismatches to infer relationships as there would be few shared SNPs among different samples. As an alternative, we calculated the correlation coefficient R of two genomes in their distance to the East Asian (ASN) samples in 1kGP, assuming that different random sampling of a fraction of the whole set of ~15K slow SNPs are roughly equivalent in representing the whole set. Verification of this R correlation approach has been described previously ^6^.

We tested the correlation of ancient samples within themselves relative to their correlation with present day Han Chinese in order to better infer the relationships among the ancient samples. We obtained for each aDNA R values with each other and with the 211 Han Chinese samples in 1kGP. We then ranked these R values as shown in Table 2, which then allowed us to infer genetic relationships among these aDNAs. We grouped these 12 aDNAs into 4 groups based on being from the same site and being closely related to each other as indicated by correlation rankings. In Table 2, rank values means the rank of a sample on the column among all 223 samples in values of correlation to a sample listed on the raw, e.g., K2 from the column was ranked 7^th^ among all samples in correlation with K12 from the raw. The 4 groups were designated as the Dashanqian (DSQ) group including the 2 samples from the Xiajiadian Culture in Dashanqian in Chifeng (K2 and K12) known to be related to the Hongshan Culture ^14^, one sample S1 from the Xiaoheyan Culture at the Hala Haigou site in Chifeng related to the Hongshan Culture ^15^ and one sample FQ17 from the East Zhou dynasty at Fushan Qiaobei Shanxi ^16^; the Banpo (BP) group including a sample BP38 from the middle Yangshao Culture site at Banpo Xian ^17^, a late Yangshao sample DZ14 from Duzhong Henan ^18^, one sample SG2046 from Sanguan Hebei known to be influenced by both Yangshao and Xiajiadian ^19^, one sample ZK22 from Zhukaigou Inner Mongolia ^20^, and one sample H69 from the early Shang site at Xuecun Henan ^21^; the Zhengzhou Xishan (ZX) group including 2 samples from the same Zhengzhou Xishan site of Miaodigou Culture in the Central Plains (ZX167 and ZX107) ^22^; and finally the Other group including PAB3002 of Gaotaishan Liaoning ^23^.

**Table 2.** Ranks in correlation between each pair of ancient samples among 211 Han Chinese in 1KG and 12 ancient samples. Rank values listed means the rank of a sample on the column among all 223 samples in values of correlation to a sample listed on the raw.

|  | FQ17 | K2 | K12 | S1 | SG2046 | DZ14 | BP38 | ZK22 | H69 | ZX167 | ZX107 | PAB3002 |
| --- | --- | --- | --- | --- | --- | --- | --- | --- | --- | --- | --- | --- |
| FQ17 |  | 170 | 173 | 169 | 220 | 126 | 182 | 152 | 78 | 74 | 207 | 221 |
| K2 | 20 |  | 7 | 90 | 217 | 206 | 209 | 61 | 169 | 69 | 167 | 193 |
| K12 | 184 | 103 |  | 5 | 222 | 180 | 198 | 174 | 109 | 190 | 192 | 143 |
| S1 | 218 | 215 | 149 |  | 218 | 188 | 206 | 204 | 106 | 211 | 193 | 196 |
| SG2046 | 221 | 219 | 222 | 151 |  | 41 | 99 | 164 | 166 | 173 | 179 | 213 |
| DZ14 | 210 | 222 | 220 | 221 | 10 |  | 1 | 21 | 5 | 215 | 219 | 215 |
| BP38 | 208 | 221 | 203 | 182 | 171 | 16 |  | 187 | 189 | 158 | 210 | 126 |
| ZK22 | 169 | 187 | 189 | 180 | 199 | 43 | 191 |  | 16 | 221 | 202 | 113 |
| H69 | 102 | 217 | 155 | 7 | 214 | 35 | 205 | 18 |  | 222 | 120 | 191 |
| ZX167 | 23 | 172 | 164 | 89 | 161 | 128 | 72 | 208 | 196 |  | 97 | 190 |
| ZX107 | 220 | 213 | 198 | 135 | 204 | 182 | 204 | 197 | 94 | 160 |  | 205 |
| PAB3002 | 222 | 214 | 151 | 61 | 215 | 136 | 90 | 123 | 133 | 204 | 178 |  |

These groupings mostly showed expected relationships based on geophysical distances, for example, the close genetic relationship of the 3 Chifeng samples in the DSQ group. Also, FQ17 in the DSQ group was located in the Y shaped region known to be a common migration route linking the Central Plains with the Northeast ^24^. A notable exception was ZX167 who had K2 and FQ17 from DSQ group among the most related aDNA samples rather than ZX107 from the same site (K2 and FQ17 ranked higher than all other aDNAs in R correlation to ZX167, being at 69^th^ and 74^th^, respectively). However, ZX107 did have ZX167 as the most related aDNA (the ranking of 97^th^ was the highest among all aDNAs in R correlation to ZX107), consistent with the geographical location.

ZX167 was the 172^th^ ranked in relation to K2 and the closest to K2 among non DSQ samples. K2 was the 69^th^ ranked in relation to ZX167 and the closest to ZX167 among all aDNAs samples here. This indicated gene flow between DSQ and ZX, which is consistent with the known archaeological findings of a migration of Miaodigou Culture to the Niuheliang site in Chifeng Inner Mongolia ^2^.

DZ14 was the first ranked among all in relation to BP38 with R values substantially higher than the next (0.38 for DZ14 and 0.24 for the second ranked) and BP38 was 16^th^ ranked among all in R to DZ14. This indicated gene flow between Banpo Shannxi (BP38) of middle Yangshao Culture and Duzhong Henan (DZ14) of late Yangshao Culture, consistent with archaeological findings.

### Relationships between aDNAs and present day samples carrying the super-grandfather haplotypes

We first determined to which present day populations the ancient Chinese samples may be closely related. We used the informative slow SNPs to calculate pairwise genetic distances between each ancient DNA and each of the 1kGP samples as described previously ^6,7^. For samples such as K2 with relatively large number of slow SNPs covered, we were able to perform informative distance analyses and principle components analysis (PCA). The results showed K2 to be closest to East Asians in genetic distances (Figure 1A). K2 was closer to Europeans than most East Asians as shown by PCA plots (Figure 1B), which is consistent with the location of K2 at Dashanqian Chifeng Inner Mongolia where East and West admixture are known to have occurred ^25^. Interestingly, K2 was more related to South East Asians such as KHV and Hunan people and least related to Northern Chinese CHB (Han Chinese in Beijing), consistent with the known migration of ancient Southwest people from Gaomiao Culture in Hunan to the Jiahu Culture in Henan and in turn to the Hongshan Culture in the Northeast as indicated by archaeological records (8 pointed star) ^26^. In contrast, PCA plots using fast SNPs clustered all aDNA samples here with PEL (Peruvian in Lima, Peru) and as outliers to the ASN samples (Figure 1C and data not shown), which was reminiscent of the surprising finding with European aDNAs that ancient samples (>2000 years old) were often not the direct ancestors of present day Europeans living in the same area ^27–30^. These results from the fast SNPs were in direct conflict with all other lines of data and hence most likely wrong, which in turn provided additional support for the theoretical justification for using slow SNPs in demographic inferences ^6–12^.

**Figure 1.**
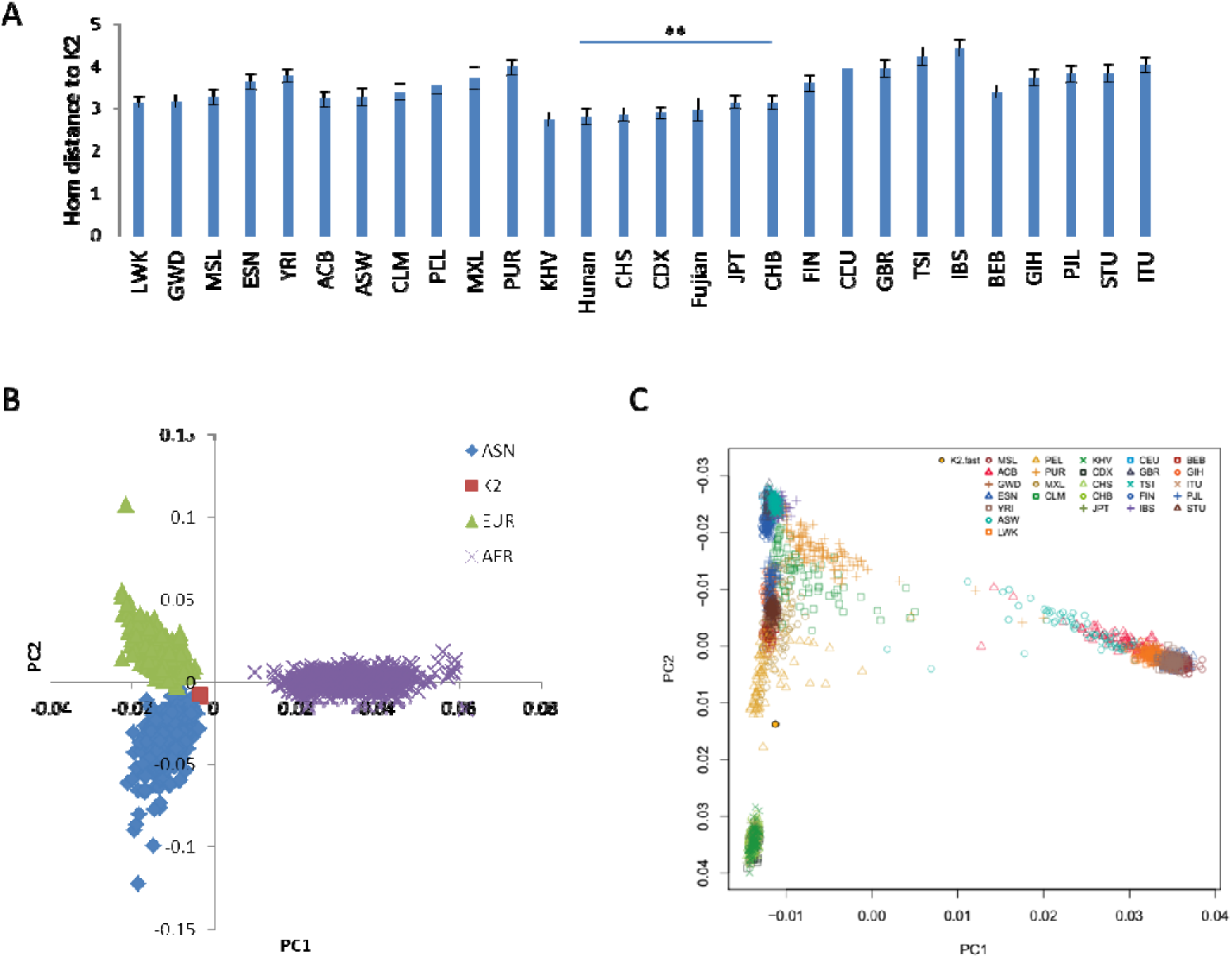
Relationship of ancient Chinese genomes with present day populations. A. Pairwise genetic distance between K2 sample and 1kGP samples. Distances were mismatches in homozygous sites. Standard error of the mean (SEM) is shown. B. PCA plots (PC1 and PC2) of K2 merged with 1kGP samples using slow SNPs. Only ASN, EUR, and AFR samples of 1kGP are selectively shown. C. PCA plots (PC1 and PC2) of K2 merged with 1kGP samples using fast SNPs.

We performed correlation analyses using distances to European samples of 1kGP to determine whether the aDNAs here were more related to East Asians (CHS Southern Han Chinese) or Europeans (CEU, Utah Residents with Northern and Western European Ancestry). All informative aDNAs with R values greater than expected by random simulations (>0.02) showed greater affinity to CHS than to CEU (Table 3). Some samples had two few SNPs to be informative. Therefore, although the numbers of SNPs here were limited, which would weaken the strength of correlations, most samples were still informative enough to be able to properly identify the correct group affiliations of these aDNAs.

**Table 3.** Correlation of ancient samples from China with Han Chinese (CHS) or Europeans (CEU) of 1kGP. Shown are Spearman correlation R values in distance to EUR samples in 1kGP. Two samples from Han Chinese of 1kGP were shown as positive controls. Comparisons were deemed non-informative when R values were 0.1 – 0 or −0.1 – 0, which happens to be found for randomly scrambled distance values as shown for CHS_HG00478.

| Sample names | R in EUR distance |  |  |
| --- | --- | --- | --- |
|  | CHS | CEU | T test |
| FQ17 | 0.010 | 0.039 | ni |
| K2 | 0.423 | 0.229 | <0.01 |
| K12 | 0.179 | 0.165 |  |
| S1 | 0.125 | 0.080 | <0.01 |
| SG2046 | 0.165 | 0.152 |  |
| DZ14 | 0.136 | 0.126 |  |
| BP38 | 0.197 | 0.126 | <0.01 |
| ZK22 | 0.182 | 0.073 | <0.01 |
| H69 | 0.158 | 0.138 | <0.05 |
| ZX167 | 0.306 | 0.211 | <0.01 |
| ZX107 | -0.024 | 0.010 | ni |
| PAB3002 | 0.114 | 0.088 | <0.01 |
| CHS_HG00478 | 0.623 | 0.376 | <0.01 |
| CHB_NA18611 | 0.512 | 0.380 | <0.01 |
| CHS_HG00478 random | -0.008 | -0.020 | ni |

Our previous studies showed that individuals carrying the same Y haplotype were also more related in autosomes, which could only be shown with slow but not fast autosomal SNPs ^7^. We next determined the autosomal relationship of each aDNA to present day people carrying the super-grandfather Y haplotypes. We calculated the average rank in R values of each aDNA to the F5 samples relative to the F11 and F46 samples in the 211 Han Chinese in 1kGP. A ratio <1 in the F5 rank versus the F11 or F46 rank means closer relationship to the F5 samples relative to the F11 or F46 samples. Relative to F11 samples, DSQ and ZX but not BP were found closer to F5 samples, indicating that the DSQ and ZX groups had more F5 associated autosomes and less F11 associated autosomes whereas the BP group had the opposite (Figure 2A). Relative to the F46 samples, DSQ and BP groups were more evenly related to F5 and F46 but the ZX group was still more related to F5, indicating that the ZX group had probably the most F5 associated autosomes among the aDNAs examined here. BP38 was the only sample in the BP group that had significantly more F11 than F46 (P <0.01, chi-squared test), indicating close relationship of BP38 with the F11 haplotype known to be common in Southern China. While DSQ and ZX groups were both related to F5, DSQ was more related to F46 than ZX.

**Figure 2.**
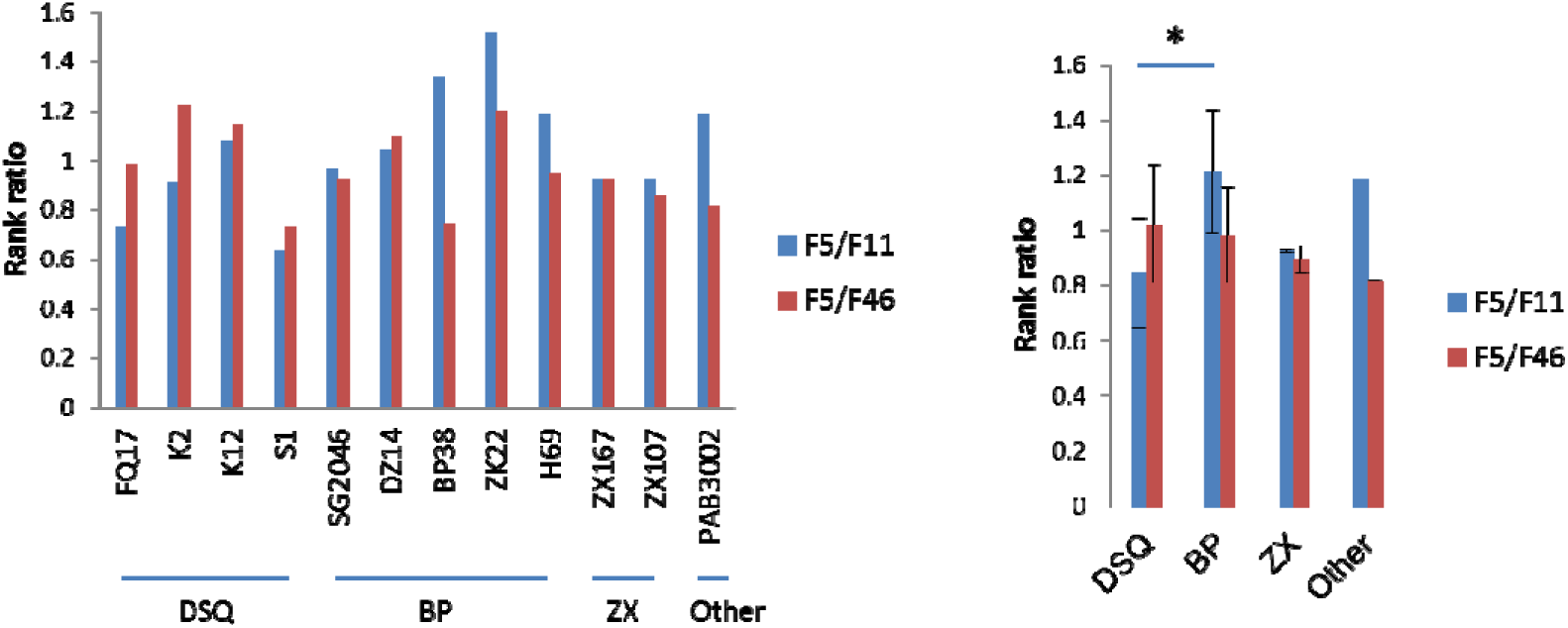
Autosomal relationships of aDNAs with Han Chinese samples of 1kGP carrying F5, F11, or F46 haplotypes. R correlation values with 211 Han Chinese and 12 aDNAs were obtained and ranked for each aDNA sample. The average rank to F5, F11, or F46 was calculated for each sample listed here and the ratios in the ranks are shown either for each sample (A) or the average of each group (B).

We also did the same analysis using total pairwise distance values calculated from the set of fast evolving SNPs, as would normally be done by the field. The results showed that all 3 aDNA groups (DSQ, DZ, and ZX) had F5/F11 ratio <1 (0.81-0.84), indicating no sensible correlation of any kind. Such dramatic difference in results obtained from the fast and the slow SNPs was consistent with previous findings ^6,7^, confirming again that only slow SNPs could be used for informative analyses in phylogenetic studies or in detecting specific association between autosomes and Y haplotypes.

To determine further the possible Y haplotypes associated with the aDNA groups, we studied the autosomal relationship of aDNAs with present day samples grouped by different haplotypes. We determined the relative distance to DSQ vs BP or to ZX vs BP for different sets of 1kGP samples with each set associated with a particular haplotype. For each sample, we calculated a rank in the subtraction value of ‘the rank to DSQ subtracting the rank to BP (DSQ-BP)’ with small values (<120 with 120 being the middle rank corresponding to the subtraction value 0) meaning higher rank in relation to DSQ relative to BP. We also similarly calculated a rank in the value of ‘the rank to ZX subtracting the rank to BP (ZX-BP)’. The samples of the M134 clade containing both F5 and F46 samples were significantly closer to ZX or DSQ than samples carrying O1 and O2 haplotypes (F5/F46 and O1/O2 in Figure 3). Relative to DSQ, samples carrying F11 and C were the closest to the BP group although not significant. The results suggest that BP group had more autosomal ancestry from Southern China as O1, O2, F11, and C are known to be common in the South.

**Figure 3.**
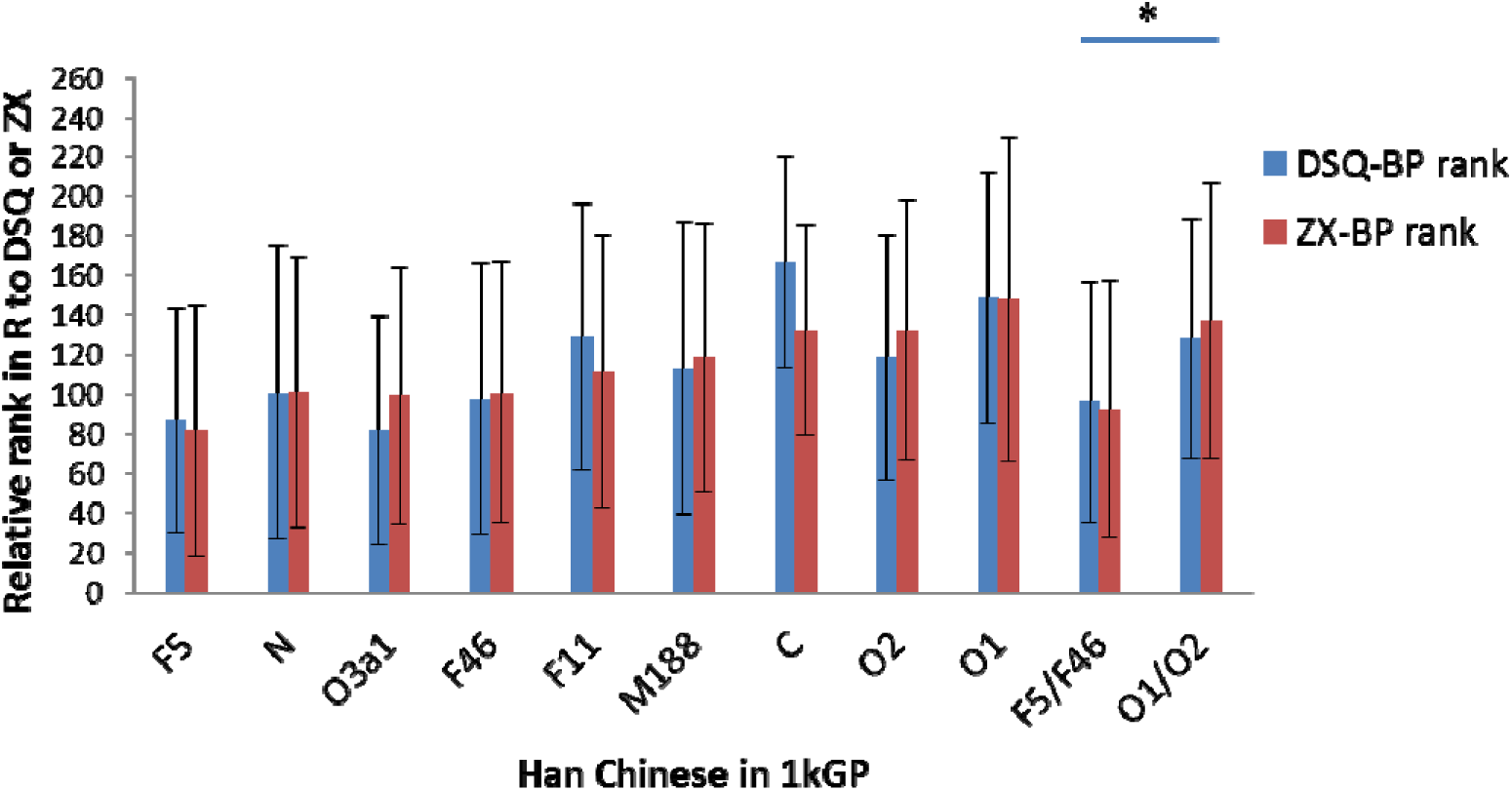
Relative autosomal distance of aDNAs to Han Chinese in 1kGP grouped by Y haplotypes. Ranks in R to aDNA groups were determined and ranks were shown for the subtraction values of ‘rank to DSQ subtracting the rank to BP (DSQ-BP)’ or ‘rank to ZX subtracting the rank to BP (ZX-BP)’. Small subtraction ranks (<120) means closer rank to DSQ or ZX, and higher value means closer rank to BP.

### Ancient DNA relationships with Southwest Chinese carrying M117 or the F5 subtype O2a2b1a1a6

Tibetans are known to commonly carry M117 ^31^ and a subtype of M117 or F5, O2a2b1a1a6, is common in Southwest Chinese ^32^. If both DSQ and ZX groups may be candidate group for the origin of F5, it would be important to ask which group is more related to Southwest people carrying M117 or O2a2b1a1a6. We made use of 4 previously published Tibetan genomes of 3150-1250 years old, including 3 M117 and one D haplotype (markers downstream of M117 were not covered) ^31^. We obtained the slow SNPs set and calculated the pairwise genetic distance with each member of 1kGP for each of the ancient Tibetans.

To determine which of the 12 aDNA samples was closely related to the M117 Tibetans, we obtained correlation R ranking of the 3 M117 Tibetans in relation to each aDNA among 227 samples including 211 Han Chinese, 12 Han aDNAs, and 4 Tibetan aDNAs (Figure 4A). The highest rank for the M117 Tibetans was found in correlation with the ZX group among the 4 aDNA groups. The results indicated possible gene flow between the ZX group and the M117 Tibetans.

**Figure 4.**
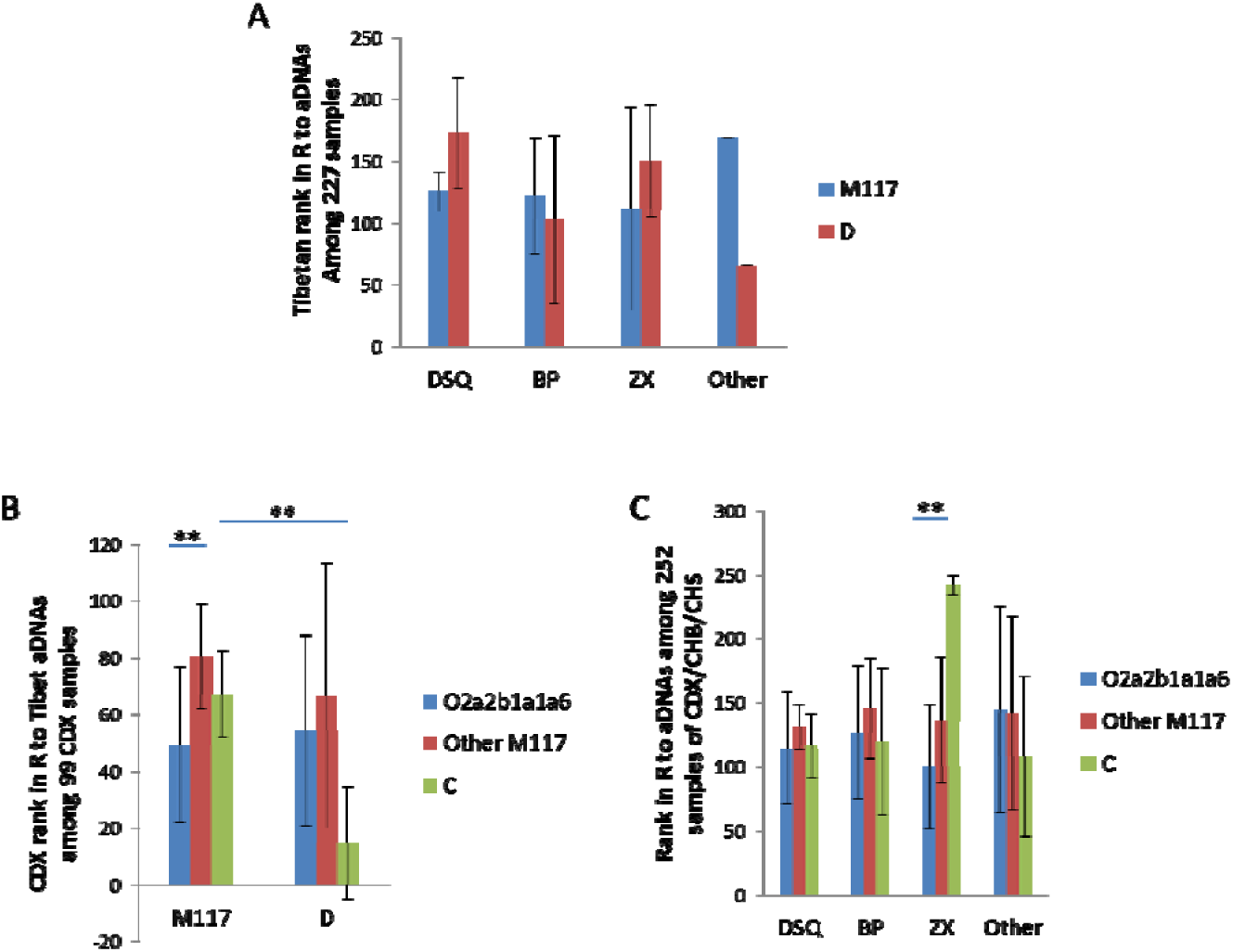
Relationship of the O2a2b1a1a6 haplotype with aDNA samples. A. Relationship between ancient Tibetans and the aDNA groups in this study. B. Relationship of Tibetans with CDX samples carrying different Y haplotypes. C. Relationship with aDNA groups of this study for CDX samples carrying different Y haplotypes.

The haplotype O2a2b1a1a6 was commonly found in Southwest Chinese such as the Dai ^32^. To further confirm the putative role of ZX in the origin of this haplotype, we studied the CDX samples (Chinese Dai in Xishuangbanna) of the 1kGP. We first confirmed that CDX samples carrying O2a2b1a1a6 were closely related to M117 Tibetans in autosomal genetic distances. We obtained R values of each aDNA with the CDX samples using their genetic distance to East Asian samples in 1kGP. We classified the M117 samples in CDX into 2 different haplogroups (O2a2b1a1a6 and other M117), and calculated the average R of the CDX in relation to the Tibetans. The results showed that CDX samples carrying O2a2b1a1a6 were significantly closer to M117 Tibetans than CDX samples carrying other M117 haplotypes. CDX samples carrying the C haplotype were significantly closer to the D type Tibetan than to M117 Tibetans, consistent with C and D belonging to the ABCDE clade ^7^. The results indicated that the autosomes of ancient M117 Tibetans were more related to those associated with O2a2b1a1a6, suggesting that ancient Tibetans likely carried the O2a2b1a1a6 haplotype.

We next divided the CDX samples into 3 Y groups (O2a2b1a1a6, Other M117, and C) and tested their relationships with the 4 groups of aDNAs here. We obtained correlation values of each aDNA to each of 99 CDX, 103 CHB, and 50 CHS samples of 1kGP. Among the 4 aDNA groups, the ZX group was found closest to the O2a2b1a1a6 samples and the only aDNA group that showed significantly closer distance to O2a2b1a1a6 samples than to C samples (Figure 4C). The results suggest a specific association of the ZX group with populations that carried the O2a2b1a1a6 haplotype, indicating either that ZX was enriched with the O2a2b1a1a6 haplotype or that ZX was the ancestral population giving rise to the group carrying the O2a2b1a1a6 haplotype. Given the present day distribution of O2a2b1a1a6 in the Southwest, it is more likely for ZX, being located in the Central Plains, to be the ancestral group to the O2a2b1a1a6 haplotype.

### Analyses of ancient Y chromosomes

Given the autosomal relationships of both the DSQ and ZX group with present day people carrying the F5 haplotype, it may be expected that at least some members of these two groups may carry the F5 haplotype. We performed Y chromosome sequence analysis on the only sample here that was good enough for high coverage sequencing analysis, the ~3000 year old K12 from Upper Xiajiabian Culture in West Liao River Valley, Northeast China ^14^. We also analyzed the published sequence of the MG48 individual from the ~4000 year old Mogou site of the Qijia Culture in Northwest China ^33^. The results showed both K12 and MG48 to be F438-F2137 with no mutations in sites that define downstream haplotypes under F2137. The results provided additional support for the presence of the F5 lineage in the DSQ group.

## Discussion

Our results here showed that aDNAs in Central and Northern China from this study could be separated into 4 groups based on autosomal relationships among them. Two among these, ZX and DSQ, were related to F5 associated autosomes. The BP group was more related to autosomes associated with the O1, O2, C, and F11 that are commonly found in the South while the ZX group was more associated with the F5 and F46 haplotypes common in the Central Plains and the North (8/12 F11 samples in Han Chinese were CHS and 6/8 F11 in CHS were from Hunan). This indicates that the BP group might be migrants from the South. Consistently, analyses of human skulls of the Duzhong and Banpo sites indicated close relationship with populations from South China ^34,35^. Human skulls from the Zhengzhou Xishan site or other Miaodigou sites such as Shanxian Henan however showed mixed features related to both Yangshao and Dawenkou people ^36,37^. Thus, people from different Miaodigou sites in Henan, such as Duzhong and Xishan, appear to have different cranial features, consistent with DNA findings here. The DNA results confirm the suggestion based on archaeological and historical records that the early Yangshao Culture and its possible predecessor the Peiligang/Jiahu Culture may be associated with migrants under the legendary *Yan* or *Fuxi* Emperor from the South such as the Pengtoushan and Gaomiao Culture ^26,38^.

The DSQ group had one sample K12 carrying F5 while the ZX group was non-informative for Y chromosome. There are at least 7 branches immediately under F5 ^32^. The F438 branch appears to be of high social economic status (SES) based on it having shorter branch length, more descendant branches, and higher fitness (lower risk for autism) ^4^. The O2a2b1a1a1a2a2 haplotype of K12 and MG48 belongs to F438. If the original F5 haplotype had a fitness advantage that might have contributed to its super-grandfather status in the first place, haplotypes with fewer random variations from the original F5 haplotype should be expected to retain the most of the fitness advantage of F5 and as such to be more likely to confer high SES status and produce more descendants. Thus populations in ancient times near the time of the original F5 would be expected to be enriched with haplotypes closest to F5. Thus the finding of two of two informative samples that have high coverage sequence data being of the F438-F2137 haplotype is consistent with *a priori* expectation of higher prevalence of a high SES haplotype in ancient times.

It appears that the ZX group may be more directly linked to the origin of F5 as it was both related to DSQ in the Northeast and the Southwest people carrying the F5 subtype O2a2b1a1a6. The common presence of F438-F2137 in the 3000-4000 YBP time period in the North indicates unlikely the presence of F* haplotype in the North in ancient times. The most parsimonious explanation for these observations is the diversification and radiation of F5 sub-branches from a centrally located population such as ZX where F* might originate.

Present day Chinese are thought to be the descendants of *Yan* and *Huang*. Based on archaeological and historical records, scholars have suspected an association of *Yan* with the Yangshao Culture in the Central Plains (but with ancestry from the South such as the Gaomiao Culture in Hunan) and *Huang* with the Hongshan Culture in the Northeast ^24,38,39^.

The Miaodigou Culture was a most popular Culture of its time and known to have impacted the Northeast HongShan Culture (and the subsequent Xiajiadian Culture to which the DSQ group belonged), more so than any other Culture of the time such as the Dawenkou Culture ^2^. Our DNA findings here suggest that there were people in the Central Plains closely related to the super-ancestor F5 lineage and possibly associated with the Miaodigou Culture. These conclusions from DNA studies are consistent with the suggestion from archaeological and historical studies that the Miaodigou Culture, and in particular the first walled town (made of rammed earth) of Xishan, may be linked to the lineage of *Huang* who is known to be the first to have built walled towns in Chinese history ^22^. People of this lineage are believed to have also lived in the Northeast (Hongshan and Xiajiadian Culture) including the great Zhuanxu Emperor, a grandson of *Huang*, and to have migrated down to the Central Plains in later times during a cold climate period ^24,39^. Samples from the Niuheliang site of Hongshan Culture are 13.7% for haplotype O2a2b1-M134 (downstream sites remain to be determined) and future studies of more samples are needed to determine if F5 was present at this site ^40^.

Overall, our study identified the presence of F5 genomes in ancient samples from the Central Plains and the Northeast and implicates the origin of the F5 lineage in the Central Plains and subsequent diversification and migration to the Northeast and Southwest. The remarkable unification of ancient DNA results with archaeological and written records can only be found when we used slow SNPs but not fast SNPs. This provides further confirmation of our new molecular methodology in demographic inferences.

## Supporting information

## Acknowledgments

This work was supported by the National Natural Science Foundation of China grant 81171880 (SH) and 31371266 (H.Z.) the National Basic Research Program of China grant 2011CB51001 (S.H.).

## Author contributions

Y.Z., X.L., and H.C. performed experiments and data analysis. H.Z and S.H. conceived and supervised the study. YZ and SH and analyzed the data and wrote the first draft of the manuscript and all authors participated in revising the manuscript.

## Materials and Methods

### Ancient DNA sequencing

Archaeological sites and samples were described in details in supplementary materials. Two teeth from each sample were collected for DNA sequencing analyses as we did in a previous study ^33^. Appropriate precautions were taken to ensure the authenticity of the ancient DNA results.

### Sequence download

We downloaded ancient and modern human genome sequences from the relevant websites using publically available accession numbers.

### Selection and identification of SNPs

Selecting SNPs: The random selection of autosomal 255K SNPs as fast evolving SNPs (none from the X chr) and the selection of the slow evolving SNPs were as described previously ^7^.

Calling SNPs: We used publically available software SAMTOOLS, GATK, and VCFTOOLS to call SNPs from either downloaded BAM files or BAM files we generated based on downloaded fastq data or our own data by using Burrows-Wheeler Aligner or BWA software ^41–43^. For better accuracy of calling SNPs, the Phred quality score (Q score) was set at 50, DP>=10, and alt frequency >=0.8. For the ancient Y chromosome DNAs, the filters for calling SNPs included DP>4 and alt frequency >=0.8.

### Genetic distance analyses and other population genetics methods

Genetic distance: Pairwise genetic distances of either hom distances (mismatch in hom sites) or total distances (mismatch in het and hom sites combined) were calculated using the custom made software “dist” as previously described ^7^. This software is freely available at https://github.com/health1987/dist and has been described in detail in previous publications ^44,45^.

### PC analysis

We utilized GCTA to analyze data in the PLINK binary PED format to generate two files (*.eigenvec and *.eigenva). We drew PCA plot using *.eigenvec file ^46,47^.

### Other methods

Other common statistical methods used were Student’s t test, chi square test, and Fisher’s exact test, 2 tailed, and Spearman correlation coefficient R analysis using Prism6 of Graphpad.

### Accession codes

The raw reads of ancient DNAs reported here have been deposited at NCBI with accession number xxxx.

