## Supplementary material for "Ancient DNAs and the Neolithic Chinese super-grandfather Y haplotypes"

**Supplementary Materials:**

**Ancient samples and archaeological sites**

S1 was excavated from the Hala Haigou site (42°21'N 11915'E) in 2007 by the archaeology institute of Inner Mongolia. This site is located in Chifeng of Inner Mongolia, and archaeologically performed the Late Neolithic Xiaoheyan culture (5000-4200 BP), which is widely distributed after Hongshan culture (6500-5000 BP) in Northeast China. Five skeletons from four different graves of Hala Haigou site were radiocarbon dated (3920±40 BP, 3850±35 BP, 3925±40 BP, 3860±35 BP, 3780±40 BP) ^1^.

K2 and K12 belonged to the Dashanqian site (41°54'N 11836'E) in Chifeng, Inner Mongolia. This site was excavated twice by the Institute of Archaeology of Chinese Academy of Social Sciences, the archaeology institute of Inner Mongolia, and Jilin University in 1996 and 1998 ^2^. Its uncovered relics showed different period cultures in Dashanqian site, including Xiaoheyan culture, Lower Xiajiadian culture, Upper Xiajiadian culture, and Zhanguo culture. Radiocarbon dates from 10 charcoal specimens from different pits and graves were analyzed by the ^14^C Laboratory of the Archaeological Science and Techniques Experiment and Research Center, IA, Chinese Academy of Social Sciences (CASS). The chronological data were 2300±50 BP, 3141±51 BP, 2451±57 BP, 2966±108 BP, 3374±55 BP, 3140±56 BP, 3180±57 BP, 3027±55 BP, 3164±57 BP, and 3184±77 BP. The selected sample K2 and K12 were found in a pit of Upper Xiajiadian culture.

PAB3002 was excavated from the Pinganbao site in 1988 by the archaeology institute of Liaoning and Jilin University. Pingganbao site is situated in Wuzhang, Liaoning province, and was used by ancient people for a very long time. Its six layers of settlements covered the periods from Late Neolithic to Bronze Age. PAB3002 was from a Bronze Age grave of the Gaotaishan culture, and was morphologically identified as a ~35 years old male ^3^.

ZK was buried in Zhukaigou site (39°6'N 1103'E) in Ordos, Inner Mongolia ^4^. This site was excavated four times in 1977, 1980, 1983, and 1984 by the archaeology institute of Inner Mongolia. A complex and characteristic culture ranging from Late Neolithic to Early Bronze Age was found in Zhukaigou site, which dated to 4200-3500 BP. It was associated with both hunting and agriculture people.

FQ17 was excavated from Qiaobei cemetery in Fushan of Shanxi province, by the institute of Qiaobei in 2003. A variety of late Shang and Zhou Dynasties style artifacts were found in all burials of this cemetery ^5^. FQ was buried in one of small graves, and was morphologically identified as a male, more than 40 years old.

SG2046 was excavated from Sanguan site in Yuxian, Hebei province by the archaeology institute of Zhangjiakou during 1979-1982 ^6^. This site shared aspects of the Lower Xiajiadian culture (2500–1500 BC), and radiocarbon dated to 3435±170 BP.^7^ Goods from this site showed successive phases from Neolithic to Early Bronze Age.

BP38 was unearthed from a four individuals multi-burial in Banpo site, which is one of the most important Neolithic sites of Yangshao culture in China. Banpo site was geographically located in Xian, Shanxi province, and had been excavated five times by Institute of Archaeology of Chinese Academy of Social Sciences during 1954-1957. A large number of remains carrying different types of Yangshao culture were found. Radiocarbon dates were obtained and cluster the Panpo site between 6700 BP and 5600 BP. BP38-4 and other three individuals were all identified as 14-15 years old females ^8^.

ZX167 and ZX107 were excavated from a Neolithic site, the Xishan site during 1993-1996. Xishan site was an old city remains of Miaodigou culture in Zhengzhou, Henan province. Its time period could trace from 6500 to 4800 years ago ^9^. ZX167 was a 35-40 years old male by morphological identification. An evidence of Tibia fracture healing was found in XS167.

DZ14 was found in Duzhong site of Mianchi, Henan province. Unearthed relics showed that this site experienced the Neolithic Late Yangshao culture and the Bronze Age Early Longshan culture. DZ14 belonged to Early Longshan culture ~4000 years ago ^10^

H69 was from Xuecun, Zhengzhou Henan and from the Early Shang period, 3500 BP ^11^.
